# CDKL5’s role in microtubule-based transport and cognitive function

**DOI:** 10.1101/2024.08.28.610038

**Authors:** André T. Lopes, Ondine Janiv, Suzanne Claxton, Sila K. Ultanir

## Abstract

Cyclin-dependent kinase like 5 (CDKL5) is a serine-threonine kinase highly enriched in mammalian neurons. CDKL5 is located on the X-chromosome and its loss-of-function leads to a severe neurodevelopmental disorder called CDKL5 deficiency disorder (CDD). CDKL5 phosphorylates microtubule-associated protein MAP1S and regulates its binding to microtubules. How MAP1S phosphorylation affects microtubule function is not well understood. To address this question, we generated MAP1S phosphomutant mice, in which the CDKL5 phosphorylation sites S786 and S812 are mutated to Alanine (MAP1S S786/812A or MAP1S SA). Using a microtubule co-sedimentation assay, we showed that dynein binding to microtubules is severely reduced in CDKL5 knockout (KO) and MAP1S SA brains. Time-lapse imaging in primary neurons showed impaired dynein motility in both Cdkl5 KO and MAP1S SA. Dynein-driven cargo transport was affected in mutant neuron dendrites, including the delivery of α-amino-3-hydroxy-5-methyl-4-isoxazolepropionic acid (AMPA) receptors. We next studied tubulin tyrosination in Cdkl5 KO and MAP1S SA neurons and found that both mutants had a reduced tubulin tyrosination when compared to WT neurons. Since dynein-dynactin has a higher affinity for tyrosinated microtubules, we hypothesized that reduced tyrosination in MAP1S phosphomutant mice could be the mechanistic cause of impaired dynein motility. In support of this, we show that upon expression of tubulin tyrosine kinase TTL, we rescued dynein motility defects in MAP1S phosphomutant neurons. Hippocampal neurons derived from MAP1S SA mice revealed a significant reduction in spine density and synapses, and altered spine morphology. Finally, behavioral phenotyping of MAP1S phosphomutant mice showed increased anxiety, impaired motor performance, social and memory deficits, mirroring to some extent the clinical manifestations present in CDD patients. Our results reveal MAP1S phosphorylation to be an important contributor to dynein-mediated transport and synapse formation.

**Graphical abstract:** 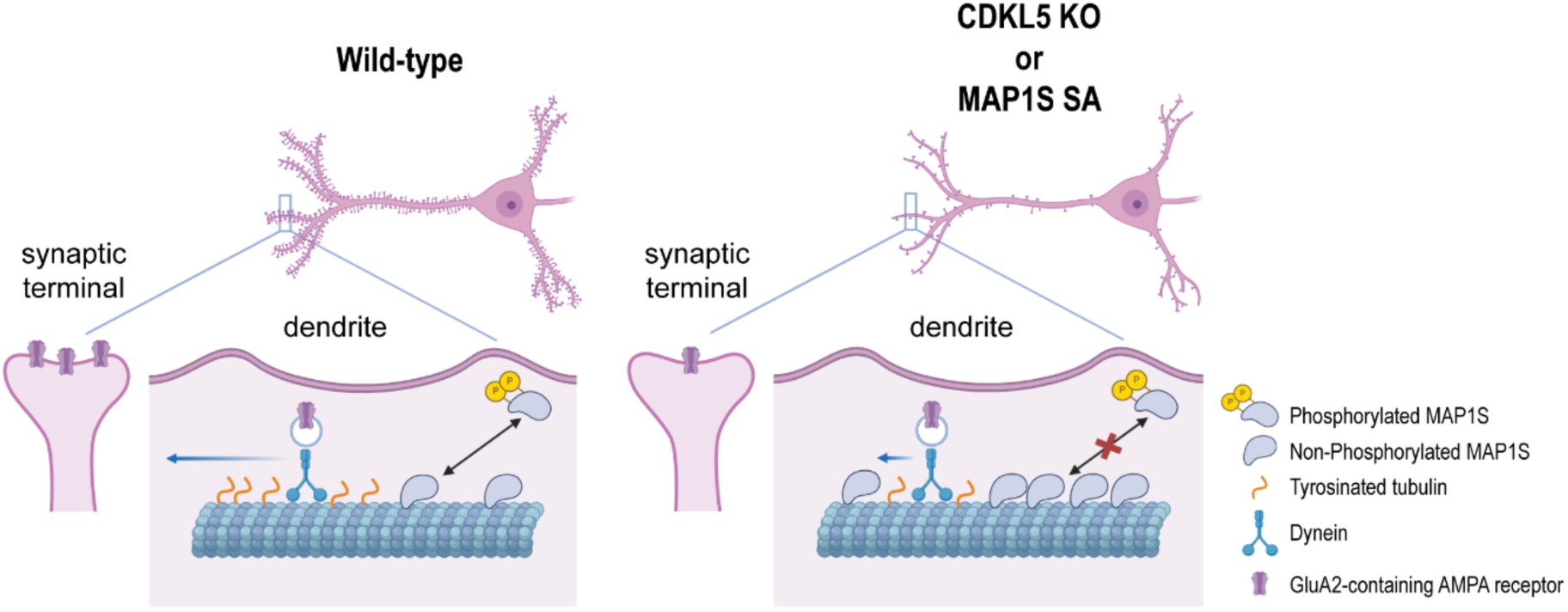

## Introduction

Cyclin dependent kinase-like 5 (CDKL5) is a 115 kDa serine/threonine kinase highly enriched in the brain, with expression starting at late embryonic development in rodents and humans^1–6^. X-linked CDKL5 mutations cause loss of CDKL5 or missense mutations in the catalytic domain, reducing the activity of this kinase and leading to CDKL5 deficiency disorder (CDD, OMIM 300203, 300672)^3,7–15^. CDD is a developmental and epileptic encephalopathy characterized by intractable early-onset seizures, severe neurodevelopmental delay, intellectual disability, and autistic features^8,16–19^. CDD is more common in females than males (at a ratio of 4:1) and affects 1 in 40,000 – 60,000 live births each year, being one of the most common types of epilepsy in childhood ^20–25^. There are currently no effective treatment options for this disorder, with CDD patients being medicated for seizures and associated comorbidities^24,26^.

CDKL5 is a kinase with an important role in the formation and maturation of neurons, cilia elongation, dendritic spine development and synaptic plasticity^27–36^. Using chemical genetics method, three microtubule-binding proteins, EB2, ARHGEF2 and MAP1S were identified as substrates of CDKL5, underscoring the importance of this kinase as a potential regulator of microtubule function^4^. Despite several studies converging on the functional link between CDKL5 and neuronal cytoskeleton^4,10,37,38^, a better understanding of CDKL5’s role in microtubule dynamics, microtubule-based transport, and cognitive function is still needed. Characterization of substrates’ roles of CDKL5 on microtubule regulation would shed light on the etiology of the disorder and lead to the development of new therapies for CDD.

Neurons depend on the microtubule cytoskeleton for numerous cellular processes such as cell division and migration, long-range transport, neurite formation and synaptogenesis^39–44^. Microtubule organization and dynamics are crucial for the cargo transport along axons and dendrites driven by molecular motor proteins like kinesins and dynein^45–50^. Microtubules are highly dynamic cytoskeletal components made of dimers of α- and β-tubulin that polymerize in a head-to-tail fashion and undergo cycles of rapid growth and disassembly in a process known as dynamic instability^51–53^. The dynamic nature of microtubules and their remodelling is necessary for neuronal intracellular transport^54^. Therefore, it is not surprising that the dynamic instability of microtubules is tightly regulated by several proteins. Microtubule-regulating proteins comprise microtubule-associated proteins, microtubule severing enzymes and enzymes performing post-translational modifications (PTMs) of α- and β-tubulin^55–58^. These PTMs not only affect the microtubule architecture but may also recruit specific effectors and provide guidance cues to the selective delivery of various cargoes such as organelles, receptors, and other signalling molecules to the correct location within the cell^59–61^.

One possible way by which CDKL5 regulates microtubule-associated functions is through the phosphorylation of its substrate MAP1S. MAP1S is a homologue of MAP1A and MAP1B and it is widely expressed in the brain. Full-length MAP1S is post-translationally cleaved into heavy chains and light chains ^62–64^. Upon phosphorylation of Serines Ser786 and Ser812, MAP1S light chain will detach from the microtubules and become more soluble in the cytosol^4^.

In this work we aimed to study the role of CDKL5-mediated phosphorylation of MAP1S on microtubule stability and organization, microtubule-based transport and animal behavior. We generated a novel MAP1S phosphomutant mouse model (MAP1S S786/812A or MAP1S SA) and used Cdkl5 KO mice^65^ to reveal the role of MAP1S phosphorylation in neurons and *in vivo*. Our results indicate that MAP1S phosphorylation by CDKL5 is essential for microtubule tyrosination, which in turn has implications on dynein-mediated dendritic cargo trafficking, synaptogenesis and ultimately mouse behavior.

## Results

### CDKL5 phosphorylation regulates dynein affinity to microtubules and dynein motility

To investigate the relevance of MAP1S phosphorylation by CDKL5 we generated a novel MAP1S phosphomutant mouse model (MAP1S S786/812A or MAP1S SA) using CRISPR- Cas9^66^. It is known that microtubule-binding proteins affect microtubule motor functions^55^. Thus, we hypothesized that MAP1S phosphorylation could influence motor protein association with microtubules. To study the role of MAP1S phosphorylation on motor protein binding, we performed a cosedimentation assay using brain lysates from Cdkl5 knockout (KO) and MAP1S phosphomutant mice (SA) (**Fig. 1a-d**). The microtubules were pelleted with the associated proteins after the incubation with the ATP analogue adenylyl-imidodiphosphate (AMP-PNP), which stabilizes motor protein-microtubule interactions^67^. The western blotting analysis indicated that the total protein levels of dynein were comparable among brain lysates from Cdkl5 KO, MAP1S SA and their respective wild-type control mice. However, there was a significant reduction of dynein affinity to microtubules in both mutant mice, when brain lysates were treated with AMP-PNP and pelleted by ultracentrifugation (**Fig. 1.c** and **d**). These data indicated that less dynein is associated with microtubules when MAP1S is not phosphorylated by CDKL5 at Ser786 and 812.

**Figure 1:**
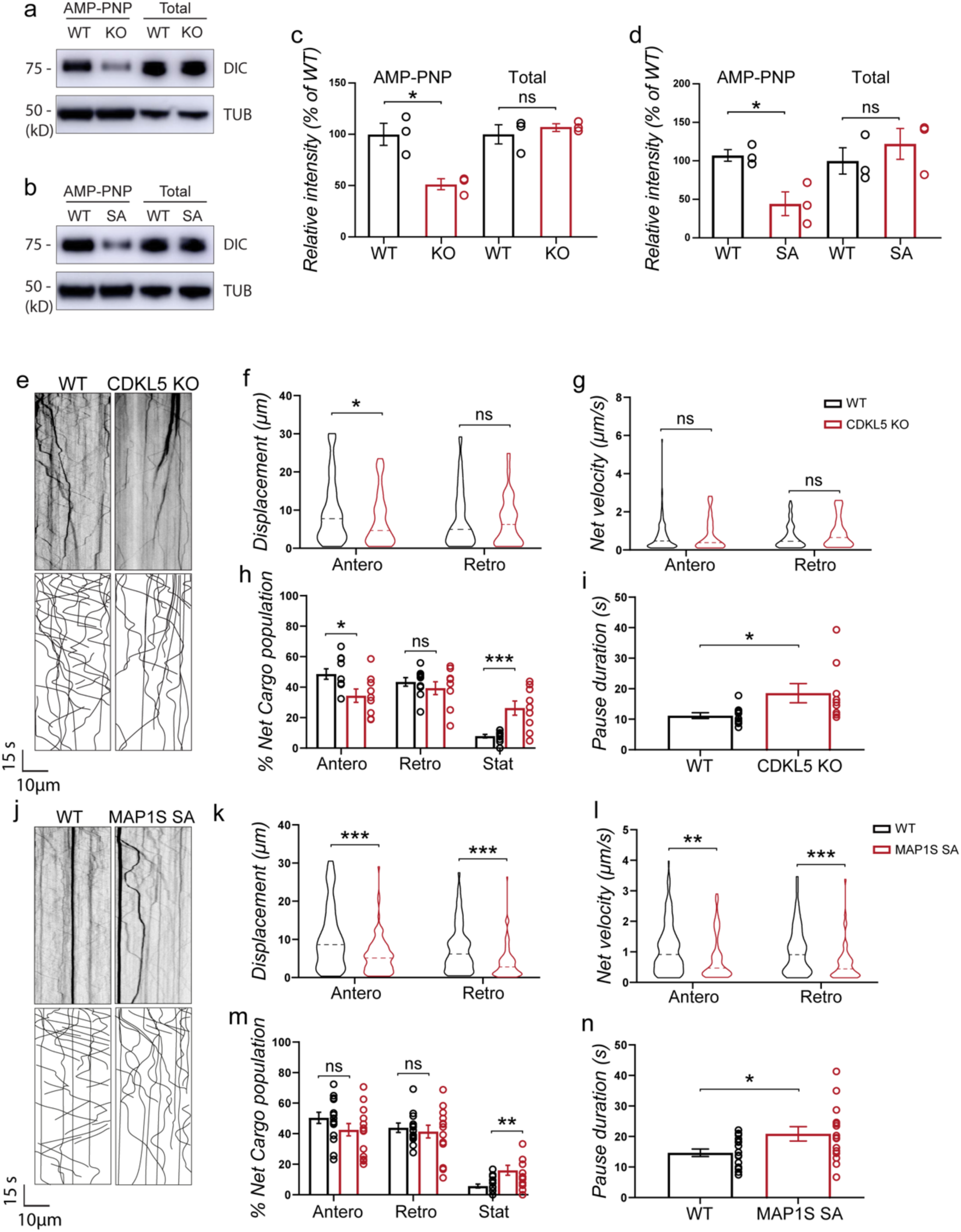
Effects of CDKL5-mediated phosphorylation of MAP1S on binding of dynein to microtubules and dynein motility. **a** and **b**) Crude microtubules were prepared from cortical homogenates of WT, Cdkl5 KO and MAP1S phosphomutants in the presence of AMP-PNP. **c** and **d**) Total and pelleted DIC (Dynein intermediate chain) were detected by western blot analysis. Signal intensities were quantified and are represented as percentage of the mutant signal relative to wild-type signal. WT, n=3, Cdkl5 KO, n=3, MAP1S SA, n=3. **e**-**n**) Live imaging of dynein-pex transport in dendrites from WT and Cdkl5 KO and MAP1S SA hippocampal neurons at DIV7. **e** and **j**) Kymographs. **f** and **k**) Displacement. **g** and **l**) Net velocity. **h** and **m**) Percentage of net cargo populations. **i** and **n**) Pause duration. Images were taken at 7 frames per second. WT, n=10-16 cells, Cdkl5 KO, n=9 cells, MAP1S SA, n=15 cells. Two-tailed Student’s *T* test was used to assess statistical significance. ns: not significant, *p<0.05, **p<0.01 ***p<0.001. Data are represented as mean±SEM.

Since dynein binding to microtubules was impaired in Cdkl5 KO and MAP1S SA, we hypothesized that neuronal transport may be altered in these mice. To test this, we took advantage of an inducible intracellular motility assay to track dynein motility in hippocampal neurons upon rapalog 1 treatment^50^. CDKL5 is highly active in dendrites as shown by phosphorylation of its target EB2^4^, therefore we focused on dendritic transport. We found that dynein motility was impaired as shown by a reduction in anterograde transport, an increase in the number of stationary particles and increased pause duration in dendrites of Cdkl5 KO neurons (**Fig. 1e-i**). Notably, these defects were even more pronounced in MAP1S phosphomutant neurons, which suggests that other kinases may compensate for the absence of CDKL5 kinase in Cdkl5 KO neurons (**Fig. 1j-n**)^68^. Taken together, these results suggest that when MAP1S phosphorylation is absent or deficient, dynein-microtubule affinity is reduced, impacting dynein-driven transport.

### Loss of MAP1S phosphorylation alters dynein-driven transport of peroxisomes and AMPA receptors

The efficient transport of proteins, RNA and organelles is crucial for neuronal function. Dynein is a large motor protein that drives transport towards the minus end of microtubules within axons and dendrites^69^. Therefore, we aimed to assess whether MAP1S phosphomutant-derived neurons display any transport deficits. We transfected primary neurons with Lamp-1-RFP to visualize the transport of lysosomes along dendrites (**Fig. 2a**). Analysis of the time-lapse imaging videos showed significantly reduced travel distances (**Fig. 2b**) and transport velocities (**Fig. 2c**) in the absence of CDKL5 phosphorylation as compared to wild-type controls. The directionality of lysosome transport did not seem to be affected in MAP1S SA neurons. There was a trend for more stationary lysosomes (**Fig. 2d**) and the duration of pauses was significantly increased in the MAP1S SA neurons (**Fig. 2e**). Next, we asked whether neurons prepared from MAP1S SA mice may be characterized by abnormal transport of synaptic cargoes like GluA2-containing AMPA receptors (**Fig. 2f**). With respect to dynein-dependent trafficking of GluA2 we observed reduced travel distances (**Fig. 2g**) and reduced retrograde velocity (**Fig. 2h**). These findings were accompanied by an increase in the percentage of stationary AMPA receptor particles (**Fig. 2i**) and longer pauses in neurons from MAP1S phosphomutant mice (**Fig. 2j**). Overall, our data shows that when CDKL5-mediated phosphorylation of MAP1S is compromised, dynein-driven transport of crucial cargoes such as lysosomes and AMPA receptors is altered in dendrites.

**Figure 2:**
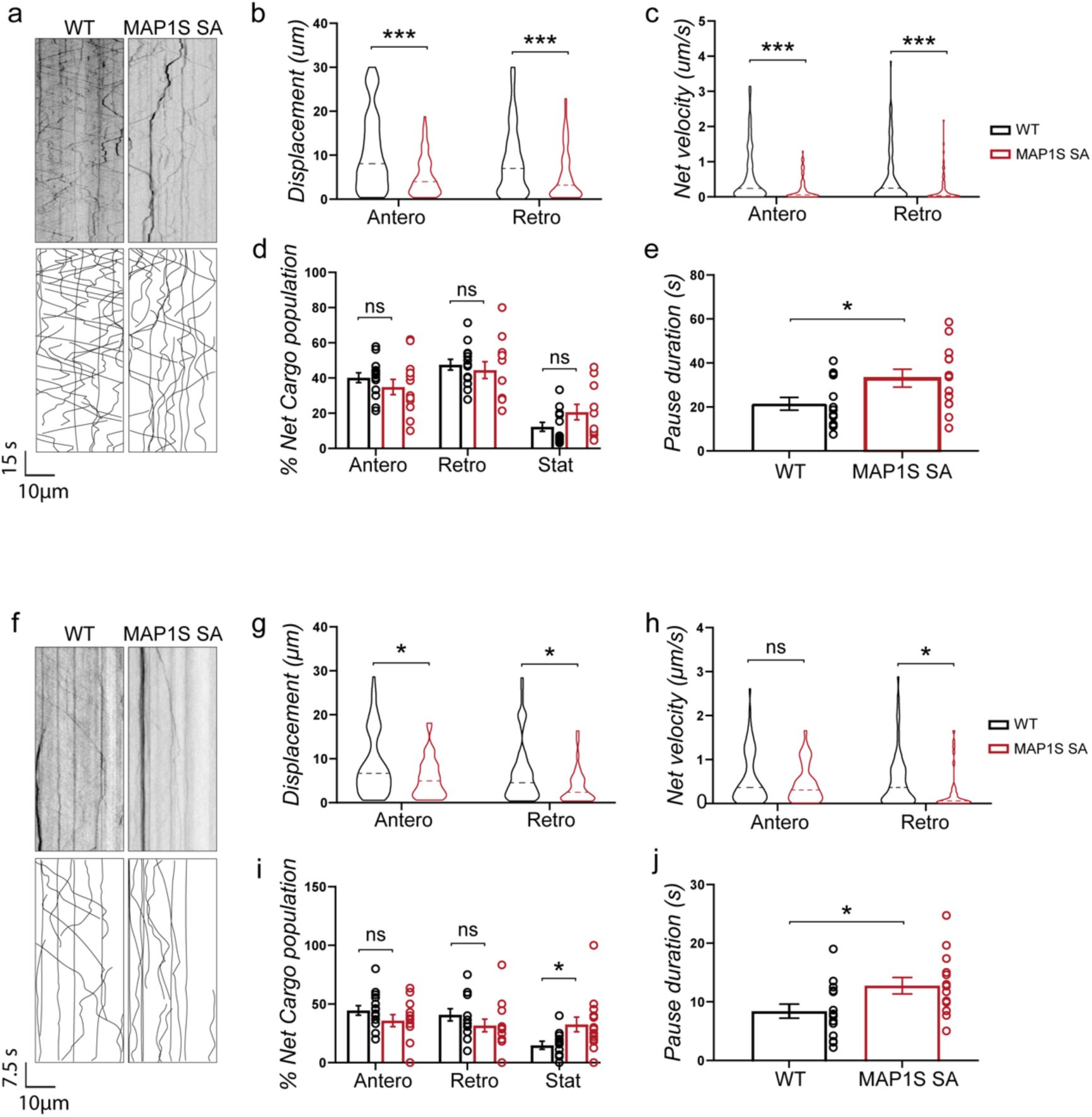
Cargo transport is impaired in MAP1S SA neurons. **a**-**e**) Live imaging of lysosome movement in dendrites from WT and MAP1S SA hippocampal neurons at DIV7. **a**) Kymographs from neurons expressing LAMP1-RFP. **b**) Displacement. **c**) Net velocity. **d**) Percentage of net cargo population. **e**) Pause duration. Images were taken at 5 frames per second. WT, n=14 cells, MAP1S SA, n=13 cells. **f**-**j**) GluA2 transport in dendrites from WT and MAP1S SA hippocampal neurons at DIV7. **g**) Displacement. **h**) Net velocity. **i**) Percentage of cargo populations. **j**) Pause duration. WT, n=15, MAP1S SA, n=14. Two-tailed Student’s *T* test was used to assess statistical significance. ns: not significant, *p<0.05, **P<0.01, ***p<0.001. Data are represented as mean±SEM.

### CDKL5 phosphorylation regulates spine morphology and synaptic densities

Since the motor protein dynein is involved in the transport of various cargoes through different neuronal compartments, we asked whether MAP1S SA neurons have an abnormal distribution of synaptic cargoes. Since dynein-driven postsynaptic GluA2 subunit-containing AMPAR transport was markedly reduced in MAP1S SA neurons we studied surface GluA2 expression levels in secondary dendrites in cultured neurons (**Fig. 3a**). Immunofluorescence staining of surface GluA2 receptors showed reduced intensity (**Fig. 3b**) and smaller GluA2 clusters in MAP1S SA neurons (**Fig. 3c**). Additionally, we biochemically analyzed total GluA2 levels (S1 fraction) and GluA2 levels in membrane-enriched fractions (P2 fractions) prepared by differential centrifugation^70^ (**Fig. 3d**). Again, MAP1S SA lysates showed lower levels of surface GluA2 receptors, (**Fig. 3f**) with total GluA2 levels remaining comparable in both groups (**Fig. 3e**).

**Figure 3:**
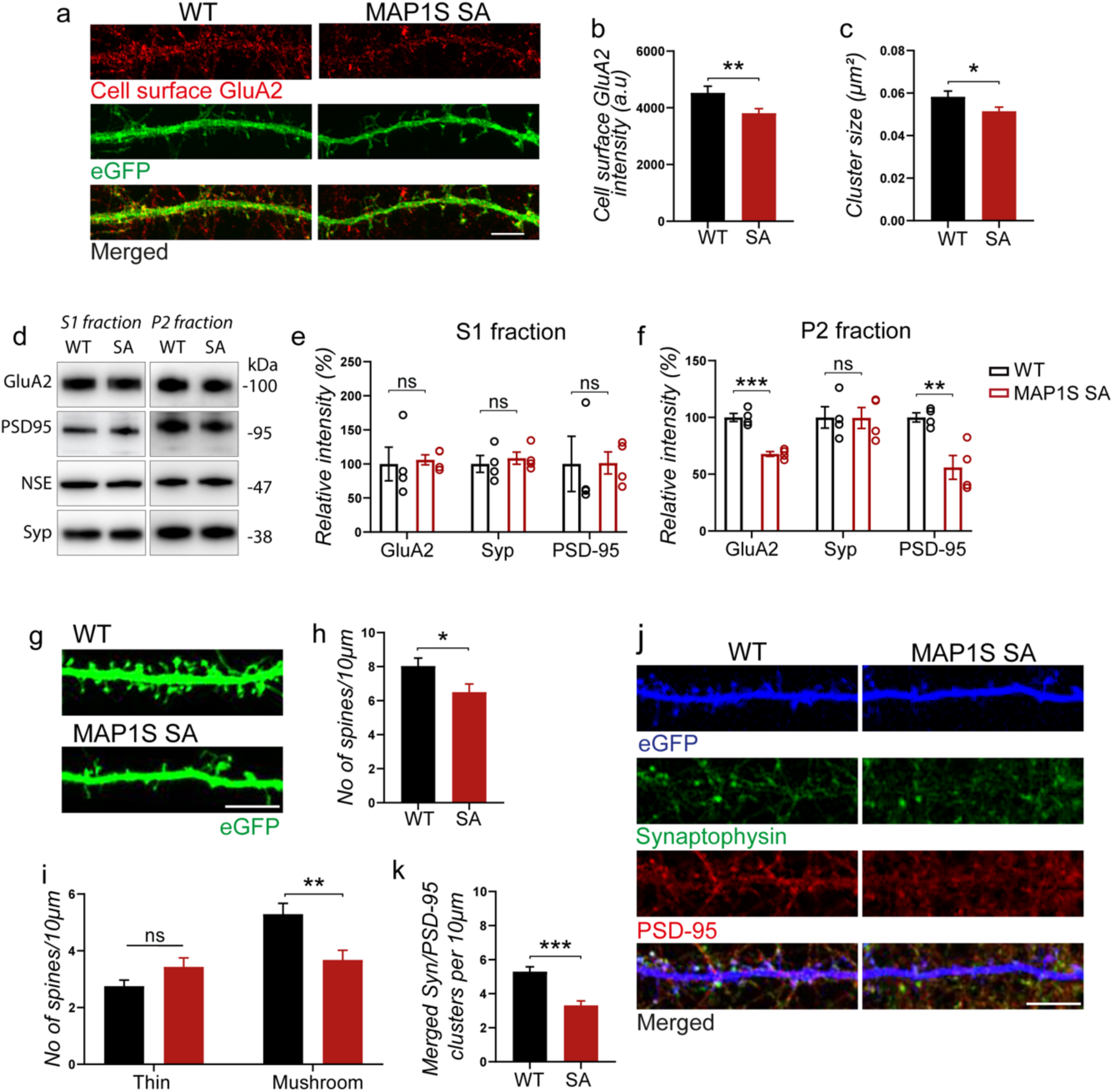
MAP1S phosphorylation regulates spine morphology and synapse density. **a**) GluA2-AMPAR cell surface staining on dendrites of hippocampal neurons from WT and MAP1S SA mice at DIV14. Cell surface (red) expression in dendrites, GFP as a volume marker. **b** and **c**) Quantification of cell surface GluA2 intensity and GluA2 cluster size, respectively. Scale: 5 μm. WT, n=16 cells, MAP1S SA, n=16 cells. **d**-**f**) Western blot analysis of GluA2, PSD-95 and synaptophysin levels normalized against NSE in S1 fraction (total levels) and P2 fraction (crude synaptosomal fraction). WT, n=4; MAP1S SA, n=4. **g**) Representative images of eGFP-transfected neurons at DIV18. Scale: 5 μm. **h**) Quantification of total number of spines per 10 μm of dendrite length. **i**) Quantification of spine types per 10 μm dendrite length. WT, n=24 cells, MAP1S SA, n=29 cells. **j**) Coimmunostaining of Syp and PSD-95 at DIV18 to visualize excitatory synaptic contacts. Scale: 5 μm. **k**) Quantification of excitatory synapses per 10 μm dendrite length at DIV18. WT, n=24 cells, MAP1S SA, n=29 cells. Two-tailed Student’s *T* test was used to assess statistical significance. ns: not significant, *p<0.05, **P<0.01.***p<0.001. Data are represented as mean±SEM.

Neuronal cells rely on proper intracellular transport carried out by various motor proteins to deliver important cargoes such as receptors into the synapse. This is not only important during the formation of synapses but also for their maintenance and normal function. Indeed, defects in intracellular transport are often concomitant with abnormal synapse morphology and physiology. Altered AMPA receptor content correlates with spine morphology^71^. We therefore performed a morphometric analysis of the dendritic spines and analyzed the number of synaptic contacts (**Fig. 3g**). In MAP1S SA mice, the spine density in secondary dendrites was significantly reduced compared to wild-type littermates (**Fig. 3h**). Interestingly, the number of mushroom-type spines was mainly affected by the loss of MAP1S phosphorylation and there was a trend for an increased density of immature spines (thin spines) (**Fig. 3i**). Next, we transfected hippocampal neurons from WT and MAP1S SA mice with eGFP and fixed them at DIV18. The neurons were further stained against the pre-synaptic protein synaptophysin and the postsynaptic protein PSD-95 (**Fig. 3J**). Colocalized pre- and postsynaptic markers, representing glutamatergic synapses, were significantly reduced in the MAP1S phosphomutant neurons as compared to wild-type neurons (**Fig. 3k**). Our data therefore suggest that CDKL5-mediated phosphorylation of MAP1S represents a critical parameter in AMPA receptor surface expression, dendritic spine maturation and excitatory synapse numbers.

### CDKL5 phosphorylation regulates microtubule stability and dynamics

MAP1S directly binds to microtubules and affects their bundling and stability^4,62,72^. To investigate microtubule stability in MAP1S SA neurons, we compared the resistance of microtubules from Cdkl5 KO and MAP1S SA hippocampal neurons to the depolymerizing drug nocodazole (**Fig. S1a-d**). When neurons were exposed to 30 μM nocodazole for 40 minutes, there was an obvious reduction in β-III tubulin intensity in WT neurons, whereas a persistent and more marked signal could be observed in Cdkl5 KO (**Fig. S1a** and **b**) and MAP1S SA neurons (**Fig. S1c** and **d**). Live cell imaging using the fluorescent microtubule +TIP binding protein 3 (EB3) further confirmed that CDKL5-mediated phosphorylation of MAP1S affects microtubule dynamics (**Fig. 4a-e**). In MAP1S SA neurons, we observed that EB3 comets grow over longer distances both anterogradely and retrogradely when compared to WT neurons (**Fig. 4b** and **c**), mimicking EB3 dynamics in Cdkl5 KO mice^73^. However, the velocity of each run is comparable in both groups (**Fig. 4d** and **e**), meaning that CDKL5 phosphorylation of MAP1S does not affect microtubule polymerization velocity. Lack of MAP1S phosphorylation, and increased MAP1S binding to microtubules therefore affects microtubule dynamics and stability.

**Figure 4:**
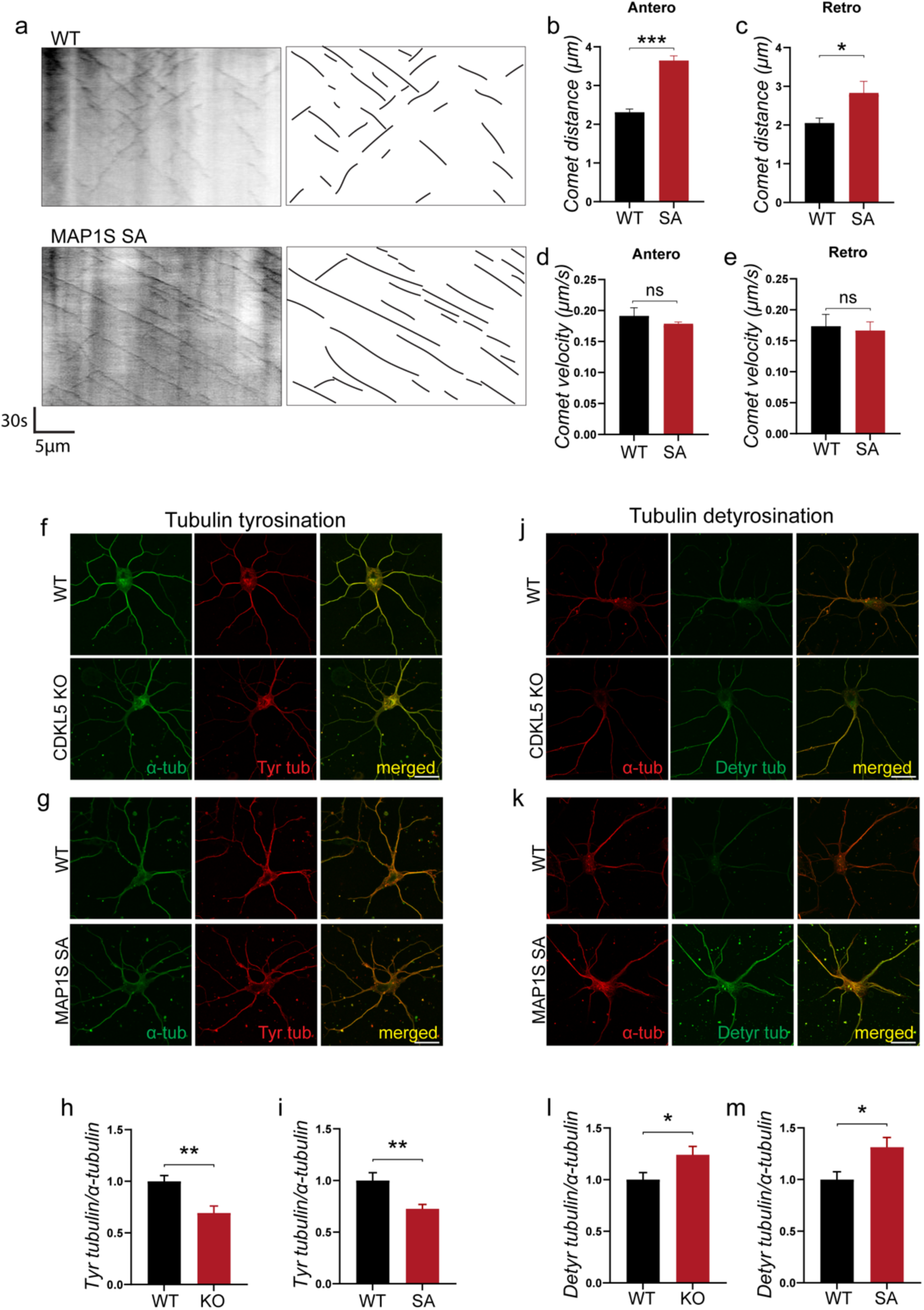
Regulation of microtubule stability and dynamics by CDKL5. **a**) Microtubule dynamics in WT and MAP1S SA neurons. Representative kymographs of EB3-tdTomato acquired in dendrites from WT and MAP1S SA neurons at DIV5. Quantification of (**b**) anterograde comet distance, (**c**) retrograde comet distance, (**d**) anterograde comet velocity, (**e**) retrograde comet velocity. WT, n=10 cells, MAP1S SA, n=11 cells. Coimunostaining of α-tubulin with tyrosinated tubulin (**f** and **g**) and detyrosinated tubulin (**j** and **k**) in WT, Cdkl5 KO and MAP1S SA neurons at DIV5. (**h** and **i**) Quantification of immunofluorescence signal in **f** and **g**. (**l** and **m**) Quantification of immunofluorescence signal in **j** and **k**. Scale: 20 μm. WT, n=16-33 cells, Cdkl5 KO, n=15-25 cells, MAP1S SA, n=26-28 cells. Two-tailed Student’s *T* test was used to assess statistical significance. *p<0.05. Data are represented as mean±SEM.

MAP1S binding to microtubules can alter microtubule posttranslational modifications (PTMs). The reversible addition of tyrosine side chains to the c-terminal tail of tubulins, known as tubulin tyrosination, is a well-established marker for labile, short-lived microtubules ^58,74^. Tyrosination is also one important regulator of the initiation of processive dynein^75^. We first asked whether microtubule tyrosination levels in hippocampal neurons were altered in Cdkl5 KO or MAP1S SA neurons. Immunostaining of hippocampal neurons with a tyrosination-specific antibody revealed significantly decreased levels of tubulin tyrosination in the dendrites of both Cdkl5 KO (**Fig. 4f** and **h**) and MAP1S SA (**Fig. 4g** and **i**) neurons when normalized to co-stained total tubulin. On the other hand, the levels of detyrosinated tubulin were increased in both Cdkl5 KO (**Fig. 4j** and **l**) and MAP1S SA neuronal dendrites (**Fig. 4k** and **m**). These data indicate an enhanced detyrosination in MAP1S SA dendrites.

### Tubulin tyrosination re-expression rescues dynein motility defects in MAP1S phosphomutant mice

To test whether a reduction in tubulin tyrosination was the underlying cause of the observed dynein motility deficits, we performed a rescue experiment by expressing human tubulin-tyrosine ligase (hTTL) enzyme in MAP1S SA neurons at DIV 5. Primary cultures of hippocampal neurons from MAP1S mice were transfected with either eGFP or hTTL-eGFP and co-transfected with HA-BICD2N-FRB-PCI-NEO, and GW1-PEX-mRFP-FKBP and imaged 24 hours later (**Fig. 5a**). Notably, TTL re-expression for 24 hours rescued or attenuated the dynein motility deficits in MAP1S SA neurons we previously reported (**Fig. 5b-e**). We also analyzed tubulin tyrosination (**Fig. 5f**) and detyrosinated levels (**Fig. 5h**) in neurons transfected with eGFP or hTTL-eGFP for 24 hours. As expected, hTTL-eGFP re-expression normalized tubulin tyrosination levels (**Fig. 5g**) and detyrosination levels in MAP1S SA neurons (**Fig. 5i**). These experiments show that increasing tubulin tyrosination in MAP1S SA neurons is sufficient to rescue dynein-mediated transport, indicating that defects in tyrosination in MAP1S SA neurons are a key factor in transport deficits.

**Figure 5:**
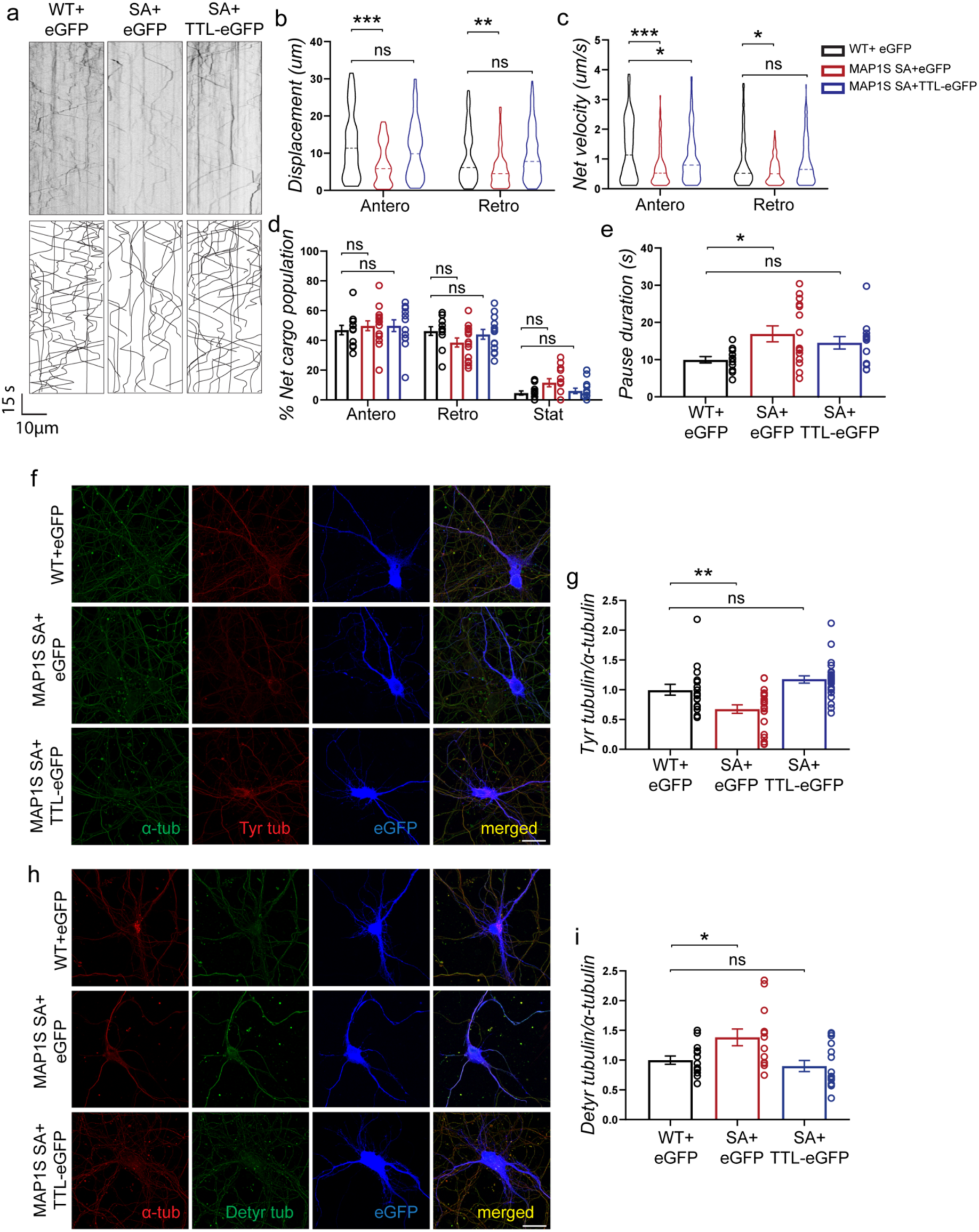
Tubulin tyrosination re-expression rescues dynein motility defects in MAP1S phosphomutant neurons. **a-e**) Live imaging of dynein-pex transport in eGFP- or TTL-eGFP transfected neurons at DIV7. **a**) Kymographs. **b**) Displacement. **c**) Net velocity. **d**) Percentage of net cargo populations. **e**) Pause duration. WT+eGFP, n=12 cells, MAP1S SA+eGFP, n=15 cells, MAP1S SA+TTL-eGFP, n=13 cells. **f** and **h**) Immunostainings for tyrosinated and detyrosinated tubulin after eGFP or TTL-eGFP expression in WT and MAP1S SA neurons. **g** and **i**) Quantification of **f** and **h**, respectively). Scale: 20 μm. WT+eGFP, n=14-18 cells, MAP1S SA+eGFP, n=13-22 cells, MAP1S SA+TTL-eGFP, n=15-26 cells. ANOVA was used to assess statistical significance. ns: not significant, *p<0.05, **P<0.01, ***p<0.001. Data are represented as mean±SEM.

### MAP1S phosphomutant mice display behavioral deficits

Several studies have reported a wide range of behavioral deficits in Cdkl5 knockout mice such as increased anxiety, poor motor performance, social and cognitive deficts^76–78^. Since these abnormalities resemble some of the clinical features of CDD patients, we were wondering whether MAP1S SA mice would show similar behavioral phenotypes. To do that, a cohort of WT and MAP1S SA mice at the age of 3 months was tested with a battery of behavioral tests. We started by assessing anxiety-like behavior in our mice. MAP1S SA mice spent less time in the open arms of the elevated plus maze (**Fig. S2a** and **b**) and less time in the center zone of the open field (**Fig. S2c** and **d**), suggesting an increase in anxiety in these mutant mice. The activity levels of MAP1S SA mice during 30 minutes of free exploration in the open field were also reduced, prompting us to assess motor performance in these mice (**Fig. S2e**). Mice were trained on the rotarod apparatus at a constant speed (4RPM) and tested 24 hours later at an accelerating speed of 4 to 40 RPM over 4 minutes (**Fig. S2f**). MAP1S SA mice had a poor motor performance shown by the reduced time spent walking on the rotarod (**Fig. S2g**). Similar results were observed for the inverted grid test, where MAP1S SA mice spent less time hanging onto the metallic grid (**Fig. S2h** and **i**). We also tested the hindlimb clasping response in MAP1S SA mice, a behavior that has been reported in mice lacking CDKL5. Unlike Cdkl5 KO mice, MAP1S SA mice do not display any hindlimb clasping (mean scores: WT mice 0.01665; MAP1S SA mice 0.1508, p=0.1151, two-tailed Student’s *T* test). To sum up, our initial behavioral assessment of MAP1S phosphomutant mice reveals increased anxiety and poor motor performance.

One of the hallmarks of CDD is the aversion to social interaction followed by the development of repetitive behaviors^79,80^. To study this behavior in our MAP1S SA, we used the three-chamber social test (**Fig. S2j**). This test takes advantage of the preference of control mice to explore a social stimulus (an unfamiliar WT mouse) over a non-social stimulus (sociability) or novel mouse over a familiar one (social novelty). During the habituation phase, in which the maze lacked both social and nonsocial stimuli, we detected no difference in activity levels between genotypes (**Fig. S2k**). In the sociability phase, both genotypes showed an overall preference for the stranger mouse over the object (**Fig. S2l**). Thereafter, in the social novelty phase, WT mice showed a clear preference for the novel mouse when compared to the familiar mouse. This was not the case for MAP1S SA mice, which showed a diminished ability to differentiate between the familiar and the novel mouse (**Fig. S2m**) and a reduction in exploratory behavior (**Fig. S2n**). Overall, MAP1S phosphomutant mice displayed social deficits, namely decreased preference for social novelty.

### MAP1S phosphorylation is important for spatial reference and associative memory

To assess the role of CDKL5-mediated phosphorylation of MAP1S in learning and memory, we employed additional behavioral tests. In a Y-maze spontaneous alternation (SA) test (**Fig. 6a**), MAP1S SA mice performed as well as the control group mice (**Fig. 6b**). These data suggest that MAP1S phosphorylation by CDKL5 is not involved in short-term spatial recognition memory. The overall exploration activity given by the total number of arm entries was comparable across all genotypes (**Fig. 6c**). We next examined context-associative conditioned fear memory in these mice using the contextual fear conditioning paradigm (**Fig. 6d** and **e**). On day 1 of the experiment, each mouse received three foot shocks and freezing behavior (absence of all movement except respiration) was monitored and used as a readout for associative fear learning and memory in rodents. WT and MAP1S SA displayed equal freezing responses during the training phase of fear conditioning, suggesting normal acquisition of the contextual fear memory (**Fig. 6f**). 24 hours later and upon re-exposure to the training context without the foot shocks (context A on day 2), MAP1S SA mice exhibited significantly lower freezing levels, pointing to problems in recalling the context-shock association (**Fig. 6g**). On the other hand, when each mouse was exposed to a different context (context B on day 3), similar freezing levels across genotypes were observed, suggesting a context-specific formation of fear memory (**Fig. 6h**). We next studied extinction of fear memory in the same cohort of mice by re-exposing them to the training context A over four consecutive days without reinforcement. As expected, WT mice showed decreased context-associated freezing levels, indicating successful weakening of the fear-learned behaviors (**Fig. 6i**). However, that was not the case for MAP1S SA mice which displayed high levels of freezing behavior over the four days. Together, these data indicate that CDKL5-mediated phosphorylation of MAP1S is critical for associative memory in mice.

**Figure 6:**
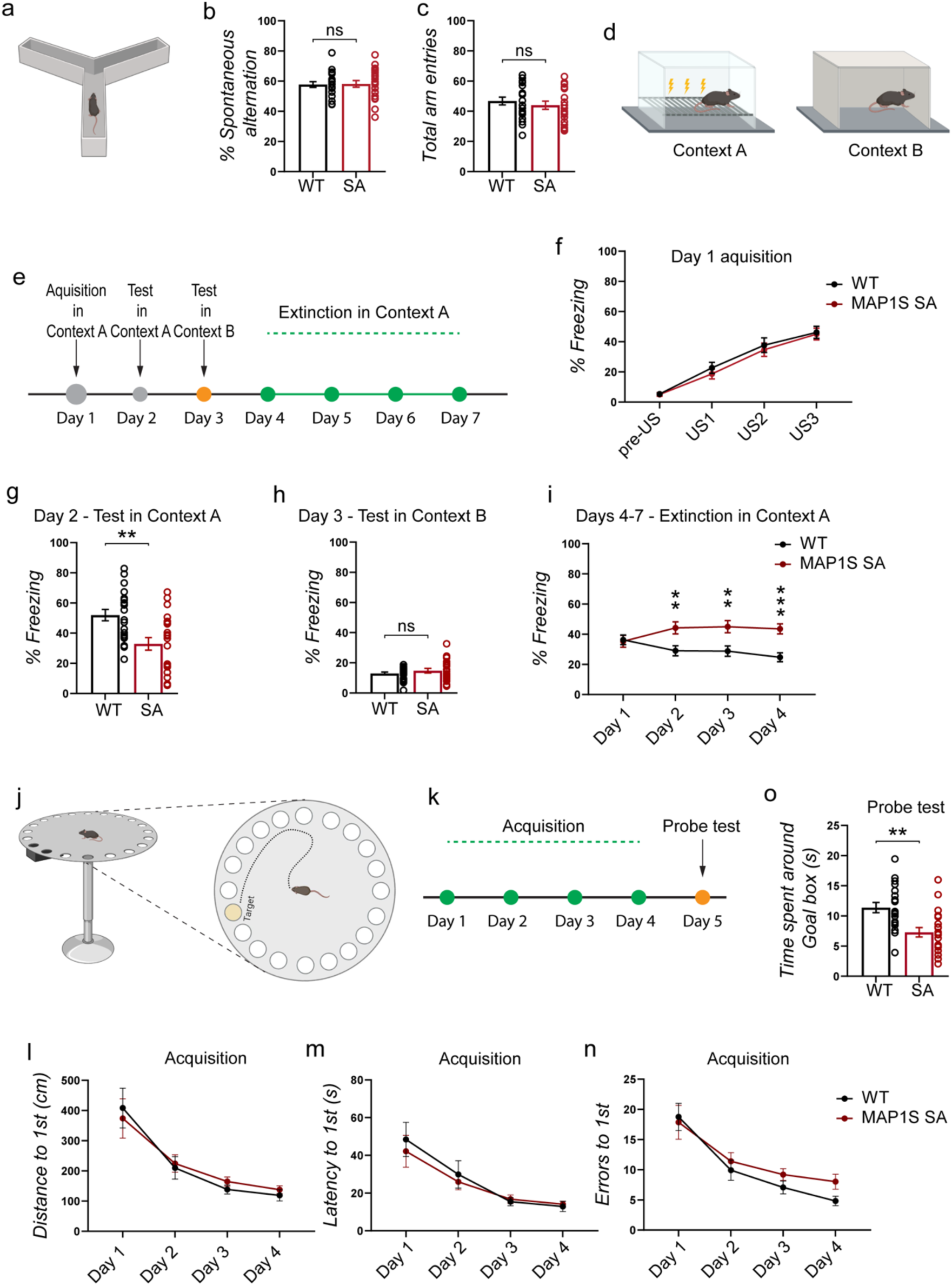
MAP1S phosphorylation regulates associative and spatial reference memory. **a**-**c**) Spontaneous alternation in the Y-maze using 3-month-old mice. **b**) Percent alternation between arm entries measures short-term spatial recognition. **c**) Explorative behavior, revealed by overall number of arm entries in the Y-maze using 3-month-old mice. **d**) Contextual fear conditioning paradigm. **e**) Experimental design of the contextual fear conditioning experiment. **f**) Day 1 acquisition in context A. Percentage of time freezing, indicative of the context/shock association. **g**) Day 2 test in context A. Percentage of time freezing, indicative of the conditioned fear memories. **h**) Percentage of time freezing at day 3, context B. **i**) Days 4-7 extinction of the fear memory in context A. Percentage of time freezing. **j**) Barnes maze test. **k**) Experimental design of Barnes maze test. **l**) Distance traveled to reach goal box. **m**) Latency to reach goal box during acquisition phase. **n**) Errors made to reach goal box. **o**) Time spent around goal box in probe test. WT males, n=10; WT females=10; MAP1S SA males, n=11; MAP1S SA females, n=10.Two-tailed Student’s *T* test, two-way ANOVA and three-way ANOVA were used to assess statistical significance. ns: not significant, *p<0.05, **P<0.01, ***p<0.001. n= Data are represented as mean±SEM. Cartoons created using BioRender.com.

We next assessed the behavior of WT and MAP1S SA mice in the Barnes maze, a test that has been widely used to study spatial learning and memory in rodents. In this test, mice are trained to find the goal box using visual cues to navigate across the maze. Cognitively intact mice spend more time exploring the goal box area or its periphery when the entry to goal box is blocked (**Fig. 6j** and **k**). Both genotypes were able to learn the task as shown by decreasing latencies, distance traveled, and errors made to reach the goal box over four consecutive days of training (**Fig. 6l-n**) even though MAP1S SA mice tended to make more errors when trying to reach the goal box (**Fig. 6n**). On the probe test day, MAP1S SA mice spent significantly less time around the goal box (**Fig. 6o**), suggesting that spatial reference memory (short-term) in these mice was impaired compared to the WT control group. Additionally, they were slower (**Fig. S2o**), made more errors (**Fig. S2p**) and traveled longer distances (**Fig. S2q**) while trying to find the goal box. Together, these results indicate that MAP1S S786/812A mice have impaired spatial reference and fear memory.

## Discussion

In this study we characterized the functional role of MAP1S phosphorylation in microtubule cytoskeleton regulation, microtubule-based transport, excitatory synapse formation and mouse behavior. Using a MAP1S S786/812A phosphomutant mouse model, we provide evidence for the critical role of MAP1S phosphorylation in neuronal synaptic development and organism function.

### Transport deficits and excitatory synapses in MAP1S SA mice

Using two mouse models, Cdkl5 KO and MAP1S SA, we showed that MAP1S phosphorylation hampers dynein affinity to microtubules and motor protein processivity. By recruiting dynein to peroxisomes using a rapalog-BicD system, we revealed a clear reduction in dynein-mediated transport in these mouse models. In addition, using RFP-tagged Lamp1 or PfusionRed-tagged GluA2, we were able to show that cargo transport was affected in dendrites. These results are in line with previous data showing a significant reduction in anterograde transport of BDNF/TrkB in the dendrites of CDKL5-depleted neurons^4^.

Reduced transport of synaptic cargo such as GluA2-containing AMPA receptors to the post-synapse may account for the abnormal dendritic spine morphology we observed in MAP1S SA mice. Spine morphological abnormalities are observed in Cdkl5 KO mice^81,82^. The trafficking of cargo along the dendrites is mainly carried out by dynein and kinesin-1 motors, and the net movement is determined by the combined activity of both motor proteins for both directions, due to the mixed polarity of microtubules in dendrites ^42,83^. Based on our data, we propose that dynein-driven transport is modulated by MAP1S phosphorylation levels, which contributes to the more modest but significant impairments of trafficking of cargos in both anterograde and retrograde directions in dendrites.

### Microtubule tyrosination as a critical modification for dynein mediated transport

In the absence of CDKL5 or when MAP1S is not phosphorylated, overstabilization of microtubules by MAP1S occurs and tubulin tyrosination is negatively regulated. In support of MAP1S’s role in microtubule stability, COS-7 cells overexpressing MAP1S light chain have very stable microtubules as shown by increased resistance to depolymerization upon colchicine treatment^62^. Additionally, our findings on Eb3 dynamics in MAP1S phosphomutant neurons recapitulate the altered microtubule dynamics reported in Cdkl5 KO neurons. In the absence of CDKL5, MAP1S has a higher affinity to microtubules and renders them less dynamic as seen by longer EB3-GFP comets, but this could be rescued by MAP1S shRNA expression^4^.

Among the microtubule PTMs, the tubulin tyrosination/detyrosination cycle has been implicated in the regulation of microtubule stability and dynamics. Briefly, when the C-terminal tyrosine of the α-tubulin is removed by a carboxypeptidase, detyrosinated microtubules are formed. The enzymatic re-addition of the tyrosine by tubulin-tyrosine ligase (TTL) leads to tyrosinated microtubules, which are more labile and dynamic^84,85^. Deficits in post-translational detyrosination/tyrosination levels have been associated with impaired axonal transport, spatial learning, synaptic plasticity and loss of synapses^86,87^.

Dynein motility impairment was due to reduced tubulin tyrosination, as rescue experiments through TTL expression, which normalized tyrosinated tubulin levels in neurons, was sufficient to rescue the dynein motility deficits in MAP1S phosphomutant neurons. It is possible that MAP1S affects the recruitment of TTL or tubulin detyrosinating enzymes (vasohibins) to microtubules^74^. For example, MAP7 association with microtubules inhibits tubulin detyrosination by sterically blocking vasohibin 1 recruitment and at the same time protects acetylated microtubules and enhances kinesin-1 binding^88,89^. We also cannot rule out the steric interference caused by increased MAP1S binding to microtubules that could render dynein binding to microtubules less efficient.

Several studies have reported that dynein and adaptor proteins like p50^glued^ and the microtubule-plus-end tracking protein CLIP-170, which enhance recruitment of dynein, bind preferentially to microtubules enriched in tyrosinated tubulin^48,55^. Importantly, CLIP-170 binding to microtubules has been reported to be deregulated in Cdkl5 KO mice. Treatment with PME (pregnenolone-methyl-ether), a compound that promotes the proper localization of CLIP-170 to microtubules was found to improve CDD-associated neuronal deficits like spine maturation and GluA2-containing AMPA receptor levels at the postsynapse^90^. These studies show that microtubule-based transport defects are a key component of CDKL5 deficiency during neuronal development.

### MAP1S phosphorylation in learning and CDD pathology

CDD is a complex clinical condition characterized by early-onset seizures, impaired locomotion and cognitive deficits accompanied by psychiatric comorbidities^91^. However, the subcellular mechanisms underlying these abnormal behaviors remain to be elucidated. In this study, we performed for the first time a comprehensive behavior analysis of MAP1S phosphomutant mice and their wild-type littermates. Our analysis revealed similarities with the previously reported deficits in anxiety, motor performance, social and cognitive impairments in mice lacking CDKL5^76,92^. However, MAP1S SA mice did not show any significant differences on the hindlimb clasping test and Y-maze. This may be explained by the involvement of other CDKL5 substrates in these behaviors in Cdkl5 KO mice. Our data support a role of MAP1S phosphorylation by CDKL5 in memory formation, most likely by controlling microtubule stability. The dynamic nature of microtubule cytoskeleton plays a crucial role in synaptic plasticity, which is widely regarded as substrate for learning and memory formation. For example, deletion of fibroblast growth factor 13 (FGF13), a microtubule stabilizing protein leads to alterations in learning and memory^93^. Another important regulator of microtubule stability is the microtubule severing enzyme spastin. Neurons lacking spastin display higher levels of polyglutamylated microtubules, reduced microtubule dynamics accompanied by cargo transport and delivery impairments, reduced numbers of excitatory synapses and cognitive deficits^94^. Interestingly, tubulin tyrosination regulates dendritic spine density, microtubule dynamics and cognition, and is downregulated in Alzheimer’s disease patients^87^.

Since CDKL5 phosphorylation promotes the unbinding of MAP1S protein from microtubules, Cdkl5 KO or MAP1S phosphomutants likely affect the cytoskeletal and synaptic rearrangements occurring during cognitive processes. However, in contrast to the memory deficits and impaired extinction of fear memory, observed in MAP1S phosphomutant mice, it is noteworthy that CDKL5-mediated phosphorylation of MAP1S does not seem to play a role in learning during the acquisition of spatial working memory in the Barnes Maze test or during the acquisition of contextual fear memory. Our study suggests that disturbing normal microtubule architecture disrupts the delivery of different intracellular cargoes, thereby reducing synaptic numbers. In particular, mushroom-type spine synapses may underlie the memory deficits found in this study.

In summary, the present study underscores the importance of MAP1S phosphorylation by CDKL5 as an essential regulator of microtubule dynamics, microtubule-based transport, and normal behavior. Our data provide a potential mechanism on how MAP1S binding to microtubules modulates microtubule architecture and composition and consequently affects dynein binding, cargo transport and cognitive function.

## Materials and Methods

### Ethics statement

All animal procedures were performed in strict accordance with the regulations of the Animal (Scientific Procedures) Act 1986 and approved by The Francis Crick Institute ethical committee. Behavioral experiments were conducted on male and female mice. The composition of the 3-month behavior cohort consisted of the following: WT males, n=10; WT females=10; MAP1S SA males, n=11; MAP1S SA females, n=10. Cdkl5 knockout mice were a kind gift from Cornelius Gross^65^.

### DNA Constructs

The following plasmids were used for transfection: EB3-tdTomato was a gift from Erik Dent (Addgene plasmid #50708), pTRIP-CAG-hTTL-eGFP and pTRIP-CAG-eGFP were a gift from Carsten Janke, HA-BICD2N-FRB-PCI-NEO and GW1-PEX-mRFP-FKBP were gifts from Lukas Kapitein, Lamp1-RFP and pCl-fusionred-GluR2(R) were a gift from Daniel Choquet. All plasmids were confirmed by DNA sequencing.

### Microtubule Binding Assay

For co-sedimentation assays, we used a modified version of a protocol previously described^67^. In brief, cortices were harvested from P20 mice and homogenized in microtubule-stabilizing buffer (1 mM EGTA/100 mM PIPES,5 pH 6.8/ 1mM MgSO4) containing protease inhibitors (10 μM PMSF/5 μg/ml E64). The cortical homogenates containing the crude tubulin were centrifuged at 50,000 x g (Beckman Coulter, Optima Max-XP) for 30 minutes at 4 °C to yield a supernatant, whereas the pellet was discarded. Afterwards, the supernatant was supplemented with 1 mM GTP, 20% glycerol, 20 μM Taxol, 1 mM AMP-PNP (pH 6.8) and incubated at 37 °C for 35 minutes, followed by a centrifugation step at 150,000 x g for 40 minutes at 37 °C. The pellet containing polymerized microtubules and co-pelleted microtubule-associated proteins was resuspended in 8 M urea for 30 minutes at RT and processed for western blotting analysis.

### Live-Cell Image acquisition

Primary hippocampal neurons were prepared as described and cultured on glass bottom dishes. Neurons were transfected using Lipofectamine® 2000 reagent protocol and imaged 24 hours later. Time-lapse live-cell imaging was performed on a Nikon CSU-W1 Ti2 Confocal microscope with a Dual Prime 95B camera with a 100x (oil) objective coupled to an incubator chamber, where hippocampal neurons were maintained in conditioned medium and kept at 37 °C and 5% CO2. For the dynein motility experiments, DIV6 neurons were co-transfected with HA-BICD2N-FRB-PCI-NEO, GW1-PEX-mRFP-FKBP and eGFP-C1 and imaged at 7 frames per second for 2 minutes. Prior to live imaging, the neurons in conditioning medium (1 ml total volume) were incubated with 8 µl of 0.1 mM rapalog 1 for 5 minutes. For the rescue experiments, in addition to HA-BICD2N-FRB-PCI-NEO, GW1-PEX-mRFP-FKBP plasmids, neurons were co-transfected with pTRIP-CAG-hTTL-eGFP or pTRIP-CAG-eGFP for 24 hours. For GluA2 transport studies, neurons at DIV5 were transfected with pCl-fusionred-GluA2(R) and imaged at 5 frames per second for one minute. To visualize EB3 comets, neurons at DIV4 were transfected with the microtubule-plus-end marker EB3-tdTomato and imaged at 1 frame per second for 3 minutes. Flat stretches of dendrites with low expression levels of EB3-tdTomato were selected for the analysis. For lysosome transport, neurons at DIV6 were transfected with Lamp1-RFP and imaged at 5 frames per second for 2 minutes. For the final analysis, captured time-lapse videos were exported as TIF images. Anterograde and retrograde movement of dynein, GluA2 single particle mobility, lysosomes as well as EB3-tdTomato comet growth of sequential images within secondary dendrites identified based on their morphology were quantified semi-manually using KymoAnalyzer plugin^95^ in Fiji (https://fiji.sc/). Background correction was achieved through subtraction of average intensity using Fiji software. Kymographs of 30 μm (Dynein, GluA2, Lysosomes and EB3 comets) dendritic segments were generated using the Multiple Kymograph plugin for Fiji.

### Nocodazole susceptibility assay

Cultured wild-type and MAP1S phosphomutant hippocampal neurons were treated with 30 μM nocodazole (Sigma) in DMSO or with DMSO alone for 40 minutes, permeabilized in PHEM buffer (60 mM PIPES, 25 mM HEPES, 10 mM EGTA and 2 mM MgCl2, pH 6.9) with 0.05% Triton^®^-X 100, and 10 μM taxol and fixed in PHEM buffer with 4% PFA. The microtubule cytoskeleton was analyzed using a beta-III-tubulin immunofluorescence.

### Behavioral Analysis

#### Y-maze spontaneous alternation test

Working memory was measured on an apparatus consisting of a Y-shaped maze with three arms made of white plastic opaque walls (34 x 9 x 14 cm) oriented in a 120° angle. For testing, the mouse was randomly placed into one arm, facing the center and allowed to freely explore the maze for 10 minutes with dim lighting (10 lux). An arm entry or exit was defined as when the mouse moved beyond the boundary of the center zone with all four paws. An actual alternation was counted when a mouse explored three different arms on overlapping triplet sets (e.g.: 1, 2, 3 or 2, 1, 3 or 3, 2, 1). The percentage of spontaneous alternation was calculated as the ratio of actual alternations to the maximum number of possible alternations (calculated as the total number of arm entries minus 2). Each trial was recorded using a video-tracking system (Ethovision XT 15).

#### Barnes-maze test

The Barnes maze consisted of a white, horizontal circular platform (96 cm in diameter, elevated 90 cm above the floor) containing 20 evenly spaced holes. Around the maze, four signs were hung on the walls of the room to serve as navigation cues. The maze was kept in a brightly illuminated room (635 lux) and underneath the target hole, a dark escape box (20 × 11 × 7 cm) was fixed. Each mouse was placed in the center of the maze and allowed to freely explore it for 3 minutes. In case the mouse did not find the escape box, the experimenter will gently guide the mouse to the target hole. Each mouse was tested in two trials per day for four consecutive days (training phase) with an inter-trial time of at least 20 minutes and the location of the escape box was kept constant. To assess learning, a 3-minute probe test was performed with the escape box being removed, and the entry blocked before the mouse was tested. Distance to first: distance traveled from the center of the arena to the target hole (center-point detection); latency to first: time to reach the target hole from the center of the arena (center-point detection); errors to first are defined as checking any hole before reaching the escape box (nose-point detection).

#### Contextual Fear Conditioning

Contextual fear conditioning is a widely used paradigm to study learning and fear memory after aversive stimulus, typically a mild foot shock is delivered. On day 1, the mouse was placed in the conditioning shock chamber (context A) consisting of a box with opaque walls (27 x 27 x 35 cm) and equipped with digital NIR video fear conditioning system (Ugo Basile, Italy) coupled to a computer. Each trial lasted 8 minutes and every 2 minutes a foot shock (unconditioned stimulus, US) was given to the mouse (0.25 mA, duration: 1 s). Contextual visual cues were added to the context A to assist with the association context-foot shock: vanilla sent, a visual sign on one of the walls and the light was kept on (100 lux). On day 2, To examine the conditioned response to the context, mice were placed back into the same context but without receiving foot shocks for 5 minutes. On day 3, mice were introduced to a different context for 5 minutes (context B), where the grid floor was covered with a plastic insert, no vanilla sent, no illumination and no visual cue was added. From day 4 to day 7, extinction of freezing behavior in mice was assessed. Fear extinction is expected to occur when context A no longer predicts unconditioned stimuli, which is described as a new form of learning). The freezing behavior of each mouse was videotaped and analyzed using Ethovision XT15 (Noldus, Leesburg, VA).

#### Quantification and statistical analysis

All quantifications throughout this study are based on at least 2-3 independent experiments. Immunoblotting was performed using standard methodologies and the quantification of optical densities of respective proteins was done using ImageJ software (NIH) or Image Studio Lite (LI-COR). Signal intensities were then normalized, as compared to loading control signal intensities and the mean of control data points was set to 100% maintaining the variance.

Maximum projection of z-stacks of confocal images (z-step: 0.5 um) was used for morphometric analysis of fixed hippocampal neurons using Metamorph software (Universal imaging). The intensity signal for each channel of the merged TIF images was determined by applying the ‘‘Integrated Morphometry Analysis’’ function and image thresholds, brightness was adjusted using the ‘‘Inclusive thresholding state’’ function.

Time-lapse images of live neurons were processed using FIJI/ImageJ. One dendrite per neuron was manually traced using the segmented line tool with a 5-pixel width and the kymograph was generated from this ROI using the MultipleKymograph plugin. The kymographs were then analyzed using the KymoAnalyzer software. For each individual track, the program calculated the net displacement, net velocity, pause duration and the percentage of particles identified as stationary or moving in the retrograde or anterograde direction. A threshold value of 350 nm was used to discriminate between stationary and mobile particles. 0.1um/s was the cutoff value set to identify a pause.

All statistical analysis and graphical representation were conducted using Graphpad Prism (version 10, Graphpad Software, Inc, San Diego, California, USA) or SPSS Statistics (version 19.0, Armonk, NY, IBM Corp). Behavioral data were first analyzed for deviation from normal normality using ‘’Shapiro-Wilk’s W’’ statistic and the homogeneity of variances using Levene’s test. Statistical significance between groups was determined by analysis of variance (ANOVA) with independent factors of genotype and sex or two-tailed unpaired Student’s *t*-test as indicated in the figure legends. When no statistical evidence for genotype x sex interactions or main effects were reported, data from both sexes were plotted together. During each experiment, the experimenter was blind to the genotype or treatment. Comparisons were considered statistically significant when *p<0.05, **p<0.01 and ***p<0.001. Data are presented as mean ± SEM.

## Supporting information

The supporting information provides further details for Supplementary Figures and Supporting Material and Methods

## Acknowledgements

We thank Erik Dent for kindly providing EB3-tdTomato (Addgene plasmid #50708), Lukas Kapitein for kindly providing the HA-BICD2N-FRB-PCI-NEO and GW1-PEX-mRFP-FKBP plasmids and Daniel Choquet for kindly providing the Lamp1-RFP and pCl-fusionred-GluR2(R) plasmids. We thank all members from Ultanir lab for their feedback and help. We thank the Crick GEMS team for the generation of MAP1S phosphomutant mice.

## Author Contributions

<u>André T. Lopes</u>, Conceptualization, Formal analysis, Investigation, Software, Data curation, Visualization, Writing; <u>Ondine Janiv</u>, Formal analysis, Investigation, Visualization; <u>Suzanne</u> <u>Claxton</u>, Formal analysis, Investigation; <u>Sila K. Ultanir</u>, Conceptualization, Data curation, Investigation, Funding acquisition, Methodology, Project administration, Supervision, Validation, Writing.

## Data and materials availability

All the data needed to evaluate the conclusions in the paper can be found in the paper and/or in the Supplementary information.

## References

1. Manning, G., Whyte, D.B., Martinez, R., Hunter, T. & Sudarsanam, S. The protein kinase complement of the human genome. Science 298, 1912–1934 (2002).

2. Chen, Q., et al. CDKL5, a protein associated with rett syndrome, regulates neuronal morphogenesis via Rac1 signaling. J Neurosci 30, 12777–12786 (2010).

3. Hector, R.D., et al. Characterisation of CDKL5 Transcript Isoforms in Human and Mouse. PLoS One 11, e0157758 (2016).

4. Baltussen, L.L., et al. Chemical genetic identification of CDKL5 substrates reveals its role in neuronal microtubule dynamics. EMBO J 37(2018).

5. Okuda, K., et al. CDKL5 controls postsynaptic localization of GluN2B-containing NMDA receptors in the hippocampus and regulates seizure susceptibility. Neurobiol Dis 106, 158–170 (2017).

6. Rusconi, L., et al. CDKL5 expression is modulated during neuronal development and its subcellular distribution is tightly regulated by the C-terminal tail. J Biol Chem 283, 30101–30111 (2008).

7. Weaving, L.S., et al. Mutations of CDKL5 cause a severe neurodevelopmental disorder with infantile spasms and mental retardation. Am J Hum Genet 75, 1079–1093 (2004).

8. Bahi-Buisson, N., et al. Recurrent mutations in the CDKL5 gene: genotype-phenotype relationships. Am J Med Genet A 158A, 1612–1619 (2012).

9. Martinez, D., Jiang, E. & Zhou, Z. Overcoming genetic and cellular complexity to study the pathophysiology of X-linked intellectual disabilities. J Neurodev Disord 16, 5 (2024).

10. Munoz, I.M., et al. Phosphoproteomic screening identifies physiological substrates of the CDKL5 kinase. EMBO J 37(2018).

11. Kalscheuer, V.M., et al. Disruption of the serine/threonine kinase 9 gene causes severe X-linked infantile spasms and mental retardation. Am J Hum Genet 72, 1401–1411 (2003).

12. Olson, H.E., et al. Cyclin-Dependent Kinase-Like 5 Deficiency Disorder: Clinical Review. Pediatr Neurol 97, 18–25 (2019).

13. Van Bergen, N.J., et al. CDKL5 deficiency disorder: molecular insights and mechanisms of pathogenicity to fast-track therapeutic development. Biochem Soc Trans 50, 1207–1224 (2022).

14. Leonard, H., et al. CDKL5 deficiency disorder: clinical features, diagnosis, and management. Lancet Neurol 21, 563–576 (2022).

15. Hector, R.D., et al. CDKL5 variants: Improving our understanding of a rare neurologic disorder. Neurol Genet 3, e200 (2017).

16. Bahi-Buisson, N., et al. Key clinical features to identify girls with CDKL5 mutations. Brain 131, 2647–2661 (2008).

17. Nemos, C., et al. Mutational spectrum of CDKL5 in early-onset encephalopathies: a study of a large collection of French patients and review of the literature. Clin Genet 76, 357–371 (2009).

18. Arts, W.F. CDKL5 gene-related epileptic encephalopathy: electroclinical findings in the first year of life. Dev Med Child Neurol 53, 296–297 (2011).

19. Fehr, S., et al. There is variability in the attainment of developmental milestones in the CDKL5 disorder. J Neurodev Disord 7, 2 (2015).

20. Symonds, J.D., et al. Incidence and phenotypes of childhood-onset genetic epilepsies: a prospective population-based national cohort. Brain 142, 2303–2318 (2019).

21. Lopez-Rivera, J.A., et al. A catalogue of new incidence estimates of monogenic neurodevelopmental disorders caused by de novo variants. Brain 143, 1099–1105 (2020).

22. Lindy, A.S., et al. Diagnostic outcomes for genetic testing of 70 genes in 8565 patients with epilepsy and neurodevelopmental disorders. Epilepsia 59, 1062–1071 (2018).

23. Kothur, K., et al. Diagnostic yield of targeted massively parallel sequencing in children with epileptic encephalopathy. Seizure 59, 132–140 (2018).

24. Bahi-Buisson, N. & Bienvenu, T. CDKL5-Related Disorders: From Clinical Description to Molecular Genetics. Mol Syndromol 2, 137–152 (2012).

25. Siri, B., et al. CDKL5 deficiency disorder in males: Five new variants and review of the literature. Eur J Paediatr Neurol 33, 9–20 (2021).

26. Jakimiec, M., Paprocka, J. & Smigiel, R. CDKL5 Deficiency Disorder-A Complex Epileptic Encephalopathy. Brain Sci 10(2020).

27. Fuchs, C., et al. Loss of CDKL5 impairs survival and dendritic growth of newborn neurons by altering AKT/GSK-3beta signaling. Neurobiol Dis 70, 53–68 (2014).

28. Zhu, Y.C. & Xiong, Z.Q. Molecular and Synaptic Bases of CDKL5 Disorder. Dev Neurobiol 79, 8–19 (2019).

29. Ricciardi, S., et al. CDKL5 ensures excitatory synapse stability by reinforcing NGL-1-PSD95 interaction in the postsynaptic compartment and is impaired in patient iPSC-derived neurons. Nat Cell Biol 14, 911–923 (2012).

30. Zhou, A., Han, S. & Zhou, Z.J. Molecular and genetic insights into an infantile epileptic encephalopathy - CDKL5 disorder. Front Biol (Beijing) 12, 1–6 (2017).

31. Katayama, S., Sueyoshi, N., Inazu, T. & Kameshita, I. Cyclin-Dependent Kinase-Like 5 (CDKL5): Possible Cellular Signalling Targets and Involvement in CDKL5 Deficiency Disorder. Neural Plast 2020, 6970190 (2020).

32. Gurgone, A., et al. mGluR5 PAMs rescue cortical and behavioural defects in a mouse model of CDKL5 deficiency disorder. Neuropsychopharmacology 48, 877–886 (2023).

33. Di Nardo, A., et al. Phenotypic characterization of Cdkl5-knockdown neurons establishes elongated cilia as a functional assay for CDKL5 Deficiency Disorder. Neurosci Res 176, 73–78 (2022).

34. Simoes de Oliveira, L., et al. Enhanced hippocampal LTP but normal NMDA receptor and AMPA receptor function in a rat model of CDKL5 deficiency disorder. Mol Autism 15, 28 (2024).

35. Awad, P.N., et al. CDKL5 sculpts functional callosal connectivity to promote cognitive flexibility. Mol Psychiatry (2023).

36. Kontaxi, C., Ivanova, D., Davenport, E.C., Kind, P.C. & Cousin, M.A. Epilepsy-Related CDKL5 Deficiency Slows Synaptic Vesicle Endocytosis in Central Nerve Terminals. J Neurosci 43, 2002–2020 (2023).

37. Barbiero, I., et al. The neurosteroid pregnenolone reverts microtubule derangement induced by the loss of a functional CDKL5-IQGAP1 complex. Hum Mol Genet 26, 3520–3530 (2017).

38. Barbiero, I., et al. CDKL5 localizes at the centrosome and midbody and is required for faithful cell division. Sci Rep 7, 6228 (2017).

39. Kapitein, L.C. & Hoogenraad, C.C. Building the Neuronal Microtubule Cytoskeleton. Neuron 87, 492–506 (2015).

40. Lasser, M., Tiber, J. & Lowery, L.A. The Role of the Microtubule Cytoskeleton in Neurodevelopmental Disorders. Front Cell Neurosci 12, 165 (2018).

41. Maday, S., Twelvetrees, A.E., Moughamian, A.J. & Holzbaur, E.L. Axonal transport: cargo-specific mechanisms of motility and regulation. Neuron 84, 292–309 (2014).

42. Hirokawa, N., Niwa, S. & Tanaka, Y. Molecular motors in neurons: transport mechanisms and roles in brain function, development, and disease. Neuron 68, 610–638 (2010).

43. Hu, X., Viesselmann, C., Nam, S., Merriam, E. & Dent, E.W. Activity-dependent dynamic microtubule invasion of dendritic spines. J Neurosci 28, 13094–13105 (2008).

44. Jaworski, J., et al. Dynamic microtubules regulate dendritic spine morphology and synaptic plasticity. Neuron 61, 85–100 (2009).

45. Miryala, C.S.J., Holland, E.D. & Dent, E.W. Contributions of microtubule dynamics and transport to presynaptic and postsynaptic functions. Mol Cell Neurosci 123, 103787 (2022).

46. Yogev, S., Cooper, R., Fetter, R., Horowitz, M. & Shen, K. Microtubule Organization Determines Axonal Transport Dynamics. Neuron 92, 449–460 (2016).

47. Guedes-Dias, P., et al. Kinesin-3 Responds to Local Microtubule Dynamics to Target Synaptic Cargo Delivery to the Presynapse. Curr Biol 29, 268–282 e268 (2019).

48. Nirschl, J.J., Magiera, M.M., Lazarus, J.E., Janke, C. & Holzbaur, E.L. alpha-Tubulin Tyrosination and CLIP-170 Phosphorylation Regulate the Initiation of Dynein-Driven Transport in Neurons. Cell Rep 14, 2637–2652 (2016).

49. Masucci, E.M., Relich, P.K., Lakadamyali, M., Ostap, E.M. & Holzbaur, E.L.F. Microtubule dynamics influence the retrograde biased motility of kinesin-4 motor teams in neuronal dendrites. Mol Biol Cell 33, ar52 (2022).

50. Kapitein, L.C., et al. Mixed microtubules steer dynein-driven cargo transport into dendrites. Curr Biol 20, 290–299 (2010).

51. Akhmanova, A. & Steinmetz, M.O. Control of microtubule organization and dynamics: two ends in the limelight. Nat Rev Mol Cell Biol 16, 711–726 (2015).

52. Akhmanova, A. & Steinmetz, M.O. Tracking the ends: a dynamic protein network controls the fate of microtubule tips. Nat Rev Mol Cell Biol 9, 309–322 (2008).

53. Gudimchuk, N.B. & McIntosh, J.R. Regulation of microtubule dynamics, mechanics and function through the growing tip. Nat Rev Mol Cell Biol 22, 777–795 (2021).

54. Kelliher, M.T., Saunders, H.A. & Wildonger, J. Microtubule control of functional architecture in neurons. Curr Opin Neurobiol 57, 39–45 (2019).

55. Goodson, H.V. & Jonasson, E.M. Microtubules and Microtubule-Associated Proteins. Cold Spring Harb Perspect Biol 10(2018).

56. Bodakuntla, S., Jijumon, A.S., Villablanca, C., Gonzalez-Billault, C. & Janke, C. Microtubule-Associated Proteins: Structuring the Cytoskeleton. Trends Cell Biol 29, 804–819 (2019).

57. Sarbanes, S.L., Zehr, E.A. & Roll-Mecak, A. Microtubule-severing enzymes. Curr Biol 32, R992–R997 (2022).

58. Westermann, S. & Weber, K. Post-translational modifications regulate microtubule function. Nat Rev Mol Cell Biol 4, 938–947 (2003).

59. Bodakuntla, S., Magiera, M.M. & Janke, C. Measuring the Impact of Tubulin Posttranslational Modifications on Axonal Transport. Methods Mol Biol 2101, 353–370 (2020).

60. Tas, R.P., et al. Differentiation between Oppositely Oriented Microtubules Controls Polarized Neuronal Transport. Neuron 96, 1264–1271.e1265 (2017).

61. Janke, C. & Magiera, M.M. The tubulin code and its role in controlling microtubule properties and functions. Nat Rev Mol Cell Biol 21, 307–326 (2020).

62. Orban-Nemeth, Z., Simader, H., Badurek, S., Trancikova, A. & Propst, F. Microtubule-associated protein 1S, a short and ubiquitously expressed member of the microtubule-associated protein 1 family. J Biol Chem 280, 2257–2265 (2005).

63. Halpain, S. & Dehmelt, L. The MAP1 family of microtubule-associated proteins. Genome Biol 7, 224 (2006).

64. Eriksson, M., et al. The NMDAR subunit NR3A interacts with microtubule-associated protein 1S in the brain. Biochem Biophys Res Commun 361, 127–132 (2007).

65. Amendola, E., et al. Mapping pathological phenotypes in a mouse model of CDKL5 disorder. PLoS One 9, e91613 (2014).

66. Hall, B., et al. Genome Editing in Mice Using CRISPR/Cas9 Technology. Curr Protoc Cell Biol 81, e57 (2018).

67. Ikegami, K., et al. Loss of alpha-tubulin polyglutamylation in ROSA22 mice is associated with abnormal targeting of KIF1A and modulated synaptic function. Proc Natl Acad Sci U S A 104, 3213–3218 (2007).

68. Silvestre, M., Dempster, K., Mihaylov, S.R., Claxton, S. & Ultanir, S.K. Cell type-specific expression, regulation and compensation of CDKL5 activity in mouse brain. Mol Psychiatry (2024).

69. Guedes-Dias, P. & Holzbaur, E.L.F. Axonal transport: Driving synaptic function. Science 366(2019).

70. Wirths, O. Preparation of Crude Synaptosomal Fractions from Mouse Brains and Spinal Cords. Bio Protoc 7, e2423 (2017).

71. Suresh, A. & Dunaevsky, A. Relationship Between Synaptic AMPAR and Spine Dynamics: Impairments in the FXS Mouse. Cereb Cortex 27, 4244–4256 (2017).

72. Tegha-Dunghu, J., et al. MAP1S controls microtubule stability throughout the cell cycle in human cells. J Cell Sci 127, 5007–5013 (2014).

73. Stepanova, T., et al. Visualization of microtubule growth in cultured neurons via the use of EB3-GFP (end-binding protein 3-green fluorescent protein). J Neurosci 23, 2655–2664 (2003).

74. Nieuwenhuis, J. & Brummelkamp, T.R. The Tubulin Detyrosination Cycle: Function and Enzymes. Trends Cell Biol 29, 80–92 (2019).

75. McKenney, R.J., Huynh, W., Vale, R.D. & Sirajuddin, M. Tyrosination of alpha-tubulin controls the initiation of processive dynein-dynactin motility. EMBO J 35, 1175–1185 (2016).

76. Okuda, K., et al. Comprehensive behavioral analysis of the Cdkl5 knockout mice revealed significant enhancement in anxiety-and fear-related behaviors and impairment in both acquisition and long-term retention of spatial reference memory. PLoS One 13, e0196587 (2018).

77. Tang, S., et al. Loss of CDKL5 in Glutamatergic Neurons Disrupts Hippocampal Microcircuitry and Leads to Memory Impairment in Mice. J Neurosci 37, 7420–7437 (2017).

78. Wang, I.T., et al. Loss of CDKL5 disrupts kinome profile and event-related potentials leading to autistic-like phenotypes in mice. Proc Natl Acad Sci U S A 109, 21516–21521 (2012).

79. Benke, T.A., et al. CDKL5 Deficiency Disorder. in GeneReviews((R)) (eds. Adam, M.P., et al.) (Seattle (WA), 1993).

80. Jhang, C.L., Huang, T.N., Hsueh, Y.P. & Liao, W. Mice lacking cyclin-dependent kinase-like 5 manifest autistic and ADHD-like behaviors. Hum Mol Genet 26, 3922–3934 (2017).

81. Della Sala, G., et al. Dendritic Spine Instability in a Mouse Model of CDKL5 Disorder Is Rescued by Insulin-like Growth Factor 1. Biol Psychiatry 80, 302–311 (2016).

82. Fuchs, C., et al. Heterozygous CDKL5 Knockout Female Mice Are a Valuable Animal Model for CDKL5 Disorder. Neural Plast 2018, 9726950 (2018).

83. Ayloo, S., Guedes-Dias, P., Ghiretti, A.E. & Holzbaur, E.L.F. Dynein efficiently navigates the dendritic cytoskeleton to drive the retrograde trafficking of BDNF/TrkB signaling endosomes. Mol Biol Cell 28, 2543–2554 (2017).

84. Sanyal, C., et al. The detyrosination/re-tyrosination cycle of tubulin and its role and dysfunction in neurons and cardiomyocytes. Semin Cell Dev Biol 137, 46–62 (2023).

85. Kreis, T.E. Microtubules containing detyrosinated tubulin are less dynamic. EMBO J 6, 2597–2606 (1987).

86. Konietzny, A., et al. Efficient axonal transport of endolysosomes relies on the balanced ratio of microtubule tyrosination and detyrosination. J Cell Sci 137(2024).

87. Peris, L., et al. Tubulin tyrosination regulates synaptic function and is disrupted in Alzheimer’s disease. Brain 145, 2486–2506 (2022).

88. Chaudhary, A.R., et al. MAP7 regulates organelle transport by recruiting kinesin-1 to microtubules. J Biol Chem 294, 10160–10171 (2019).

89. Shen, Y. & Ori-McKenney, K.M. Microtubule-associated protein MAP7 promotes tubulin posttranslational modifications and cargo transport to enable osmotic adaptation. Dev Cell 59, 1553–1570 e1557 (2024).

90. Barbiero, I., Bianchi, M. & Kilstrup-Nielsen, C. Therapeutic potential of pregnenolone and pregnenolone methyl ether on depressive and CDKL5 deficiency disorders: Focus on microtubule targeting. J Neuroendocrinol 34, e13033 (2022).

91. Dell’Isola, G.B., et al. CDKL5 deficiency-related neurodevelopmental disorders: a multi-center cohort study in Italy. J Neurol (2024).

92. Terzic, B., et al. Temporal manipulation of Cdkl5 reveals essential postdevelopmental functions and reversible CDKL5 deficiency disorder-related deficits. J Clin Invest 131(2021).

93. Wu, Q.F., et al. Fibroblast growth factor 13 is a microtubule-stabilizing protein regulating neuronal polarization and migration. Cell 149, 1549–1564 (2012).

94. Lopes, A.T., et al. Spastin depletion increases tubulin polyglutamylation and impairs kinesin-mediated neuronal transport, leading to working and associative memory deficits. PLoS Biol 18, e3000820 (2020).

95. Neumann, S., Chassefeyre, R., Campbell, G.E. & Encalada, S.E. KymoAnalyzer: a software tool for the quantitative analysis of intracellular transport in neurons. Traffic 18, 71–88 (2017).

