## Supporting information for "CDKL5’s role in microtubule-based transport and cognitive function"

**Table of content**

•	Supplementary figures

•	Supporting Material and Methods

**Figure S1, related to Figure 4. Impaired microtubule disassembly in Cdkl5 KO and MAP1S SA neurons upon nocodazole treatment.** (**a** and **c**) Immunostainings for beta-III tubulin in Cdkl5 KO and MAP1S SA neurons treated with DMSO or 30µM Nocodazole for 40 minutes. **b** and **d**) Quantification of beta-III tubulin immunofluorescence signal of **a** and **c**, respectively. Scale bar: 20µm. WT DMSO, n=16; WT NOCO, n=15-20; Cdkl5 KO DMSO, n=18; Cdkl5 KO NOCO, n=18, MAP1S SA DMSO, n=17; MAP1S SA NOCO, n=14. ANOVA was used to assess statistical significance. ns: not significant, ***p*<0.01, ****p*<0.001. Data represented as mean ± SEM.

**Figure S2, related to Figure 6. MAP1S phosphomutant mice display behavioral deficits**. **a** and **b**) Elevated plus maze test. **b**) Percentage of time in open arms over 5 minutes. **c**-**e**) Open field test. **d**) Time spent in the center zone (CZ) over 30 minutes. **e**) Total distance traveled (cm). **f-g**) Rotarod test. **g**) Graph shows latency to fall at accelerating speed over 4 minutes. **h-i**) Inverted grid test. **i**) Impulse was calculated as body weight in grams times latency to fall from the metallic grid. **j**) Experimental design of the three-chamber sociability test. **k**) Activity levels of WT and MAP1S phosphomutant mice during habituation phase as distance traveled (cm). **l**) Exploration time spent exploring object and stranger mouse during social preference phase. **m**) Exploration time of the familiar mouse versus novel mouse during the preference for social novelty phase. **n**) Time spent exploring both familiar and novel mice. **o-q**) Probe test of Barnes maze test. **o**) Latency to first (s). **p**) Errors to first. **q**) Distance to first (cm). WT males, n=10; WT females=10; MAP1S SA males, n=11; MAP1S SA females, n=10. Two-tailed Student’s *T* test and two-way ANOVA were used to assess statistical significance. ns: not significant, *p<0.05, **P<0.01, ***p<0.001. Data are represented as mean±SEM. Cartoons created with BioRender.com.

**Supporting Materials and Methods**

**Generation of MAP1S phosphomutant mice.** Mutant MAP1S Ser786/812A animals were generated in-house as part of the Francis Crick Institute’s Genetic Modification Services. Briefly, MAP1S S786A and S812A mice were generated by the cytoplasmic microinjection of Cas9 mRNA, in vitro transcribed paired guide RNAs (gRNAs), sgRNA1 (GGCAGCTACAGTTGCAGCTC), sgRNA2 (GGCTCACTGCGGGCGCTGAG), and a 200bp single stranded oligonucleotide repair template (CTCAACACCACCAATCCCAGCCGTAGCCGCAAGGCCCCTGCAAGGCCTAGCgCTGCCTCTGCCACCCCtAGAGCTGCAACTGTAGCTGCCAAGACAAAGGGCCCCGCGGGcGAtCGGAACCGGCCACTCgcCGCCCGCAGTGAGCCTGCAGACAGGCCAGGCCGTGTGCCCCTTACCAGGAAGCCCTCAGTCCCTAAGAC) into a fertilized single cell staged C57BL/6J embryos and then surgically transferred into pseudo pregnant CD1 foster mice. Genotype was determined using Transnetyx based Q-PCR probe assays and then further validated if targeted using short amplicon deep Miseq Illumina Sequencing. Wild-type and MAP1S SA mice were generated from heterozygous matings and housed in an animal facility at the Francis Crick Institute under a reversed 12-hour light/dark cycle. Temperature (20-24 °C) and humidity (55±10%) in the animal facility were kept constant, and the animals had ad libitum access to food and water. The behavioral experiments were conducted during the dark (active) phase of the cycle.

**Primary Neuronal Culture of Hippocampal Neurons and Transfection.** Primary cultures of hippocampal neurons were prepared from embryonic day 16.5 mice. Hippocampi were isolated and enzymatically dissociated with 0.05% trypsin/EDTA for 5 minutes at 37°C (Invitrogen, Waltham, MA, USA) followed by blocking with DMEM:F12 (Gibco) containing 10% Fetal bovine serum. Hippocampi were mechanically dissociated in HEPES buffer (10 mM HEPES, 135 mM NaCl, 5 mM KCl, 2 mM CaCl2, 2 mM MgCl2, 15 mM Glucose; pH 7.4). Then, 60,000 neurons were plated on 12 mm glass coverslips pre-coated with Poly-L-Lysine (50 μg/ml) (Sigma-Aldrich, Buchs, Switzerland) and kept in an incubator at 37 °C and 5% CO2. Hippocampal neurons were transfected using lipofectamine 2000. Briefly, 2 μg plasmid DNA was diluted in 200 μl of Neurobasal medium (Gibco, 21103-049), containing 6 μl Lipofectamine 2000 reagent (Invitrogen, 11668-019). The mixture was incubated at room temperature (RT) for 20 minutes and the neuronal cells were incubated with the DNA solution for 15 minutes at 37ºC with 5% CO2. Hippocampal neurons were cultured in the original (conditioned) medium for indicated time points at 37ºC and 5% CO2.

**Antibody Application Details and Reagents.** The following antibodies were used for immunofluorescence or western blotting analysis: mouse anti-α-tubulin (Abcam, AB7291, ICC, 1:500, WB 1/5000), mouse anti-tubulin-βIII (Biolegend, 801201, ICC 1:500, WB 1/10 000), rabbit anti-detyrosinated (Millipore, AB3201, ICC 1:500, WB 1:10000), mouse anti-tyrosinated tubulin (Sigma, T9028, 1:10000), mouse anti-PSD95 (Thermo Fisher Scientific, MA1-046, ICC 1/2000, WB 1:500), guinea pig anti-synaptophysin-1 (Synaptic Systems, 101004, ICC 1/2000, WB 1:500), mouse anti-GluA2 (Millipore, MAB397, ICC 1/300, WB 1/500), chicken anti-neuronal-specific enolase (Novus Biologicals, NB100-046, WB 1/5000), mouse HRP-conjugated (Jackson , 715-035-151, WB 1/10 000), rabbit HRP conjugated (Jackson, 711-035-152, WB 1/10 000), chicken HRP-conjugated (Dianova, 103-035-155, WB 1/2500), mouse AF568 (Life technologies, A10037, ICC 1/500), rabbit AF488 (Jackson, 711-545-152, ICC 1/500), guinea pig Cy5 (Jackson, 706-175-148, ICC 1/500). Nuclei were counterstained with DAPI (Merck, Darmstadt, Germany; 124653, ICC 1/1000). Reagents used include nocodazole, taxol (sigma) and rapalog 1 (Ariad Pharmaceuticals).

**Protein Extraction and Western Blot Analysis.** For the western blots, cortices, and hippocampi from WT and MAP1S SA mice were dissected out and placed in 1.5 ml ice-cold homogenization buffer (20 mM HEPES, 100 mM potassium acetate, 40 mM KCl, 5 mM EGTA, 5 mM MgCl2, pH 7.2, protease inhibitor (Roche Complete, Millipore, 11873580001), 2 mM mg ATP, 5 mM DTT and 1 mM PMSF). The tissue was then homogenized by vigorous shearing with 5 passages through a 1 mL syringe with a 21-gauge needle and 10 passages with a 26-gauge needle. Samples were then spun at 1000 x g for 10 minutes at 4°C (Thermo Scientific Heraeus Fresco 21) to yield S1 and P1 fractions. Part of the resulting supernatant (S1) containing the total protein fraction was centrifuged for 10 minutes at 10,000 x g (Thermo Scientific Heraeus Fresco 21). The supernatant was transferred into a new tube (S2) and the pellet (P2) containing plasma membrane-enriched fraction was resuspended in homogenization buffer. The S1 and P2 samples were then mixed in 4 x sample buffer containing 400 mM DTT, boiled at 95 °C for 10 minutes, and stored at -80 °C. Ten micrograms of total protein were subjected to SDS-PAGE and transferred to a PVDF membrane, which was then blocked in 5% milk in tris-buffered saline containing 0.1% Tween-20 (TBST). Primary antibodies were added for 60 minutes at room temperature or overnight at 4 °C and after washing, the membrane was incubated for 60 minutes with the HRP-conjugated secondary antibody. Bands were visualized by a chemiluminescence detection system (Amersham Imager 600 RGB). The acquired images were processed using Adobe Photoshop 2024 and the intensities of individual bands were measured using the ImageJ software (ImageJ 1.8).

**Immunostainings.** For all the confocal microscopy analysis, cultured hippocampal neurons on glass coverslips were fixed with 4% paraformaldehyde/4% sucrose diluted in PBS for 10 minutes. After fixation, cells were washed three times in PBS, followed by 4 minutes of permeabilization with PBS containing 0.5% triton-X-100, and blocked in PBS containing 1% (w/v) BSA for an hour at room temperature. Neurons were then incubated for one hour in primary antibodies diluted in blocking buffer at RT or overnight at 4°C. After a wash step, secondary antibodies and DAPI were added for one hour. Coverslips were mounted in Aqua-Poly/Mount. (PolySciences, 18606-20). This protocol was followed for all the standard immunocytochemistry, except for surface GluA2 staining, when neurons were incubated with the primary antibody under nonpermeabilizing conditions for 2 hours at room temperature followed by fixation and permeabilization for 4 minutes and blocking with 1% (w/v) BSA in PBS.

**Behavioral analysis**

**Elevated plus maze.** Animals were placed in an apparatus consisting of two open arms (5 cm x 30 cm) and two closed arms (5 cm x 30 cm with 16 cm tall opaque walls) with a center area of 5 cm x 5 cm and elevated 80 cm above the floor. The maze was assembled in a quiet and dimly illuminated room (70 lux). Briefly, each mouse was placed in the central zone facing an open arm and allowed to explore the maze for 5 minutes. Movement and time spent in the closed arms was recorded by a video camera coupled to a computer running the Ethovision tracking system (version XT 15, Noldus Technology, Wageningen, The Netherlands).

**Open field test.** The mouse was placed in the center of an opaque arena (50 x 50 x 40 cm) made of PVC foam material with a white floor and brightly lit by an overhead light (70 lux). The center of the maze, a 16 cm x 16 cm square was pre-defined as the central zone. Mouse tracking was automated using Ethovision tracking system over 30 minutes (version XT 15, Noldus Technology, Wageningen, The Netherlands). Total distance traveled and percentage of time spent in the central zone were used as measures for locomotor activity and anxiety levels, respectively.

**Hindlimb clasping test.** This test was used to assess motor deficits and is indicative of disease progression in mouse models of neurodegeneration. Mice were suspended by the base of the tail at least 15 cm above the ground and their behaviors were recorded for up to 15 seconds. Hindlimb clasping was rated from 0 to 3 based on severity: 0 = if the hindlimbs were consistently splayed outward and away from the abdomen, 1 = if one hindlimb was retracted towards the abdomen for at least 50% of the time, 2 = if both hindlimbs were partially retracted for at least 50% of time, 3 = if both hindlimbs were completely retracted and touching abdomen for at least 50% of the observation period.

**Inverted grid suspension test.** This test was used to study grip strength and coordination in mice. Each mouse was placed on a square metallic grid with the edges covered, which was slowly turned over. The grid was hold in position 40 cm above a padded surface in a brightly illuminated room (70 lux). Each experiment was stopped after the mouse fell off or after 300 seconds spent hanging from the metallic grid. The mouse was videotaped using a fixed video camera (Sony HDR-CX240E, Tokyo, Japan) and the performance was evaluated by measuring the latency to fall.

**Accelerating Rotarod**. A standard rotating rod (Ugo Basile model 47650, Italy) was used to assess balance and motor coordination at accelerating speed. Soft tissue was placed below the rotating rod for the mouse to land on. Briefly, 24 hours before the experiment, each mouse was trained for two trials with a inter trial interval of an hour, to walk on a rotating rod at constant speed (4 RPM) for three minutes. Next day, the mouse was placed on the rotating rod and the rotation speed gradually increased from 4 to 40 RPM within four minutes. A second and third trial were conducted one hour after the first trial. A video camera (Sony HDR-CX240E, Tokyo, Japan) was used to record each trial and performance was evaluated by measuring the latency to fall.

**Three-chamber social interaction test.** The apparatus was an open-topped box made of PVC foam (62 × 42 × 23 cm) divided into three chambers of equal size connected by square openings. Empty cylindrical wire cage (diameter: 8 cm, height: 18 cm) was placed in the center of each lateral chamber. During the habituation phase, each of the chambers contained an empty cylindrical wire cage and test mouse was allowed to freely explore all three chambers for 10 minutes. After this, the mouse was kept in a holding cage for the next three minutes. During sociability phase, an unfamiliar mouse (stranger mouse) of the same sex, age, strain and weight was enclosed in one of the cylindrical wire cages. The location of the stranger mouse was alternated between the two side chambers across test mice. In the other wire cage, an unfamiliar and inmate object was introduced as non-social stimulus. The test mouse was placed in central chamber of the apparatus containing the social and non-social stimuli for 10 min and then, returned to the holding cage for additional three minutes. During the social novelty phase, a new unfamiliar mouse (novel mouse) was added to the wire cage that had contained the object during the sociability phase and the stranger mouse is used as familiar one. The test mouse was placed in the central chamber and allowed to explore the arena for the next 10min, after which it was returned to the home cage. Exploration of the social and non-social stimuli was defined as when the test mouse ’s nose was detected within 2 cm from the cylindrical wire cage. A video camera positioned above the arena was coupled to a computer running Ethovision software (Ethovision XT 15) for automated mouse tracking.
